# A dynamical model of TGF-*β* activation in asthmatic airways

**DOI:** 10.1101/2021.03.28.437380

**Authors:** Hannah J. Pybus, Reuben D. O’Dea, Bindi S. Brook

## Abstract

Excessive activation of the regulatory cytokine transforming growth factor *β* (TGF-*β*) via contraction of airway smooth muscle (ASM) is associated with the development of asthma. In this study, we develop an ordinary differential equation model that describes the change in density of the key airway wall constituents, ASM and extracellular matrix (ECM), and their interplay with subcellular signalling pathways leading to the activation of TGF-*β*. We identify bistable parameter regimes where there are two positive steady states, corresponding to either reduced or elevated TGF-*β* concentration, with the latter leading additionally to increased ASM and ECM density. We associate the former with a healthy homeostatic state and the latter with a diseased (asthmatic) state. We demonstrate that external stimuli, inducing TGF-*β* activation via ASM contraction (mimicking an asthmatic exacerbation), can perturb the system irreversibly from the healthy state to the diseased one. We show that the properties of the stimuli, such as their frequency or strength, and the clearance of surplus active TGF-*β*, are important in determining the long-term dynamics and the development of disease. Finally we demonstrate the utility of this model in investigating temporal responses to bronchial thermoplasty, a therapeutic intervention in which ASM is ablated by applying thermal energy to the airway wall. The model predicts the parameter-dependent threshold damage required to obtain irreversible reduction in ASM content suggesting that certain asthma phenotypes are more likely to benefit from this intervention.

## 1 Introduction

The onset of asthma remains poorly understood. Asthma is a chronic lung disease characterised by inflammation, airway hyperresponsiveness and airway remodelling (Brightling et al., 2012; Berair et al., 2013). Inflammation refers to the infiltration of inflammatory cells, in response to inhaled allergen, into the airway that remain for longer than needed. These cells interact with resident cells in the airway causing production of contractile agonist which triggers ASM contraction (Kostenis and Ulven, 2006). Airway hyperresponsiveness is defined by exaggerated bronchoconstriction (narrowing of the airway) due to contraction of airway smooth muscle (ASM) in response to a relatively low dose of contractile agonist (King et al., 1999; West et al., 2013). Airway remodelling comprises the sustained structural changes due to inflammatory injury repair, airway thickening and scarring (Bossé et al., 2008; Al Alawi et al., 2014). Until recently, chronic inflammation was thought to be the main contributor to airway remodelling (Saglani and Lloyd, 2015). However, recent experimental evidence suggests that airway remodelling may be promoted by bronchoconstriction-induced airway narrowing (Grainge et al., 2011).

Bronchoconstriction activates the regulatory cytokine, transforming growth factor *β* (TGF-*β*) (Wipff et al., 2007; Buscemi et al., 2011; Tatler et al., 2011), which may further stimulate ASM contraction (Ojiaku et al., 2018), ASM proliferation (Chen and Khalil, 2006), and deposition of extracellular matrix (ECM) (Pohlers et al., 2009), thereby altering airway mechanics (Makinde et al., 2007; Burgess et al., 2016). In the healthy airway, regulatory mechanisms terminate this process (Hinz, 2015), thereby maintaining homeostasis. We hypothesise that accumulation of ASM and/or ECM up-regulates TGF-*β* production through a positive mechanotransductive feedback loop resulting in a loss of the healthy homoeostatic state. In this paper we develop an ODE-based model that accounts for the above biological processes to test this hypothesis theoretically, and to develop an understanding of the effect of multiple broncho-constrictive exacerbations on cell-ECM-mediated activation of TGF-*β* and temporal dynamics of the airway constituents.

The spindle-shaped ASM cells form bundles that surround the lumen of the bronchi (Kuo et al., 2003; Ijpma et al., 2017) and are primarily responsible for contraction, proliferation, cytokine production and ECM secretion. ASM cells express fewer contractile proteins whilst proliferating or synthesising ECM or cytokines and therefore generate less contractile force than non-proliferating ASM cells (Bentley and Hershenson, 2008). Hence, the primary roles of each phenotype differ: the contractile phenotype initiates contraction and the proliferative phenotype increases ASM mass. The ECM is a delicate mesh of deposited connective tissue and fibrous proteins (Cheng et al., 2016), that surrounds ASM bundles and provides physical support and stability to the lung (Burgess et al., 2016; Khan, 2016). Freyer et al. (2001) report that the ECM constituents fibronectin, laminin and collagen appear to aid the survival of the embedded ASM, which in turn secrete compounds to replenish the ECM to form a positive feedback loop. However, the overall accumulation of ECM does not significantly change over time within the normal healthy airway as the ECM appears to exhibit a balance between its deposition and degradation (Makinde et al., 2007).

Initially identified in human platelets as a protein associated with signalling in wound healing (Assoian et al., 1983), the pro-inflammatory cytokine TGF-*β* plays a vital role in regulating the immune system (Letterio and Roberts, 1998). TGF-*β* is secreted as part of the large latent complex (LLC) that resides in the ECM (Tatler et al., 2011). Cell-mediated stress, in response to cytoskeletal reorganisation or reaction to a contractile trigger, induces conformational changes in the LLC leading to the activation of TGF-*β* (Keski-Oja et al., 2004; Wipff et al., 2007). Herein, as in Gleizes et al. (1997) and Annes et al. (2003), we use the term ‘TGF-*β* activation’ to denote the separation of TGF-*β* from the LLC and ‘latent TGF-*β*’ indicates its inactive storage. Both the contractile ASM and the ECM are required for the mechanical activation of TGF-*β* (Tatler and Jenkins, 2012, 2015; Tatler et al., 2016). Once activated, TGF-*β* binds to cell surface receptors (Derynck and Zhang, 2003; Shi et al., 2011; Aschner and Downey, 2016) and initiates multiple cellular responses including cell contraction, apoptosis, differentiation, migration, proliferation, survival and ECM secretion (Khalil et al., 2005; Perng et al., 2006; Berair et al., 2013; Ojiaku et al., 2018).

TGF-*β* expression is increased within the asthmatic airway (Ojiaku et al., 2018) and correlates with asthma severity (Chen and Khalil, 2006; Al Alawi et al., 2014). Continual exposure to an allergen causes TGF-*β* to stimulate a dysregulated repair process of inflammation-induced injury to the airway by promoting ECM deposition and ASM proliferation which consequently thickens the airway wall (Al Alawi et al., 2014) and increases TGF-*β* secretion (Amrani and Panettieri, 2003). In addition, TGF-*β*-mediated ASM contraction may further up-regulate TGF-*β* activation. In light of these findings, it is desirable to investigate the feedback mechanisms responsible for excessive activation of TGF-*β* during an asthmatic exacerbation.

Experiments in vitro are typically reductionist in nature. To understand interactions of complex biochemical and mechanical processes requires both integrative, predictive mathematical models and experimental investigations. There are various such mathematical models that account for cell signalling events based upon experimental data (e.g. Haberichter et al., 2002; Brumen et al., 2005; Croisier et al., 2013), the majority of which employ the law of mass action to obtain ordinary differential equations (ODEs) that describe the rates of change in cytosolic and sarcoplasmic calcium concentrations that drive acto-myosin binding that in turn generates contractile force in the ASM cell. Other modelling approaches focus on acto-myosin dynamics responsible for ASM contractile force generation (Huxley, 1974; Mijailovich et al., 2000; Brook and Jensen, 2014; Brook, 2014; Rampadarath and Donovan, 2018), dynamics of transmembrane integrin connections to the ECM (Irons et al., 2018, 2020), and the transmission of contractile forces generated within ASM cells to the airway tissue. (e.g. Bates and Lauzon, 2005; Bates et al., 2009; Hiorns et al., 2014). These studies highlight the importance of ASM cell response to external mechanical loading and interactions with ECM. For instance the discrete stochastic-elastic, and related continuum, model of Irons et al. (2018) reveals that depending on the oscillatory load, two stable regimes exist where either binding or rupture events dominate, which could affect the transmission of strain generated by the contracted ASM to the airway tissue. A region of bi-stability exists between these two modes, whereby the level of adhesion depends on the loading history. In particular, Irons et al. (2018) show that ECM stiffness influences the region of bi-stability and modifies the magnitudes of the stable adhesion states. These findings could have implications for activation of TGF-*β* but this has not yet been investigated in this context.

Chernyavsky et al. (2014) present an ODE model representing ASM phenotype switching between a proliferative and contractile state. In the proliferative state, ASM is assumed to grow logistically which implicitly incorporates spatial constraints. They consider the occurrence of both periodic and of irregular exacerbations that follow a random Poisson process, in order to investigate the range of possible outcomes an individual may experience, given average airway characteristics. They find that the likelihood of developing moderately to severely remodelled airways is critically dependent on the inflammatory resolution rate and the individual’s history of induced remodelling events, and that accumulation of successive triggers have a long term impact on the degree of ASM proliferation. Similarly, the morphoelastic model of Hill et al. (2018) explores airway remodelling in response to transient inflammatory or contractile agonist challenges. Consistent with the findings of Chernyavsky et al. (2014), they find that when the inflammation resolution is slow, a small number of exacerbation events may lead to significant and persistent airway remodelling. Their results suggest that the sustained contractility of the ASM observed in asthmatics may be due to either a mechanotransductive feedback loop, insufficient clearance of contractile agonists, or a combination of the two. Building on the studies of Brook et al. (2010), Hiorns et al. (2014) and Hill et al. (2018), in our recent study (Pybus et al., 2021), we developed a biomechanical model of a lung-slice to quantify the mechanical stress experienced by the airway wall constituents in response to cyclic stretching (mimicking lung-slice stretching protocols investigating the mechanical activation of TGF-*β*). Employing both fully-3D finite element simulations and asymptotic reduction techniques, we predicted the stresses and deformation of an airway in the lung-slice, exposing the importance of the slice geometry, imposed stretch and muscle cell contractility.

Although the preceding studies couple acto-myosin, integrin or biochemical signalling dynamics to ASM cell or airway mechanics, the feedback mechanisms linking ASM and ECM interactions with TGF-*β* activation in asthma have not yet been investigated in detail. In this study, we develop an ODE model accounting for these processes to study the influence of ASM and/or ECM accumulation during airway remodelling on TGF-*β* activation. We represent phenomenologically both the mechanics associated with the ASM-ECM interaction, and the longer term effects of mechanotransduction on cell transcription processes, so as to focus primarily on the temporal dynamics of the airway constituents and of TGF-*β*. We expose a region of bistability in the model parameter space, in which both healthy homeostasis and chronic disease coexist; the unstable branch between the two acts as a transition threshold leading to irreversible alteration in airway consituents and hence development of asthma.

The development of the cited models, and the present one, aim to obtain a greater understanding of asthma pathophysiology, with a view to investigating potential asthma therapies. However, most therapeutic interventions are pharmacological, targeted at either inflammation (inhaled corticosteroids) or at ASM contraction (inhaled bronchodilators), so these models would need modification to account for relevant signalling pathways. Bronchial thermoplasty, on the other hand, involves the ablation of ASM within the airway by application of thermal energy to the airway via a bronchoscope-inserted catheter (Wilhelm and Chipps, 2016), with a consequent reduction in ASM that our model can straightforwardly replicate. The premise of this recently-developed intervention is that reduced ASM leads to reduced bronchoconstriction (Laxmanan and Hogarth, 2015; Zuyderduyn et al., 2008; Janssen, 2012; Pretolani et al., 2014, 2017; Donovan et al., 2018); however, a recent modelling-experimental-clinical study suggests that insufficient ASM reduction may be achieved for this to be the key mechanism underlying efficacy of the treatment (Chernyavsky et al., 2018). Here, we employ our model to study remodelling and TGF-*β* activation in response to bronchial thermoplasty, and in particular to understand the long term consequences of removing different amounts of ASM. Numerical simulations reveal a threshold in ASM ablation, over which a return to the homoeostatic is possible and hence airway remodelling is reversed; importantly the clinical feasibility of this threshold is dependent on the disease state, as reflected by the model parameters.

This manuscript is organised as follows; first we develop an ODE model describing the rate of change of airway wall constituents in response to TGF-*β* activation (Section 2). We then present an asymptotic reduction of this model (Section 3), that guides our subsequent fully nonlinear stability analysis (Section 4). This is followed by numerical time-course simulations to investigate the effects of external stimuli on TGF-*β* activation and accumulation of airway wall constituents (Section 5.1), and to study bronchial thermoplasty (Section 5.2). Finally, the discussion in Section 6 summarises our study.

## 2 Model development

As summarised above, active TGF-*β* (*a*^*\**^(*t*^*\**^)) binds to receptors expressed by proliferating ASM (*p*^*\**^(*t*^*\**^)), contractile ASM (*c*^*\**^(*t*^*\**^)), and other surrounding cells which initiates multiple cellular responses (Derynck and Zhang, 2003; Al Alawi et al., 2014; Aschner and Downey, 2016). These responses include the triggering of ASM proliferation and stimulation of ASM contraction.

The schematic diagram in Fig. 1 illustrates the interactions between the key airway wall components, ASM and ECM (*m*^*\**^(*t*^*\**^)), and the effects of TGF-*β* activation accommodated by our model. We incorporate density-dependent removal of active TGF-*β* (due to TGF-*β*-receptor binding events). We assume that the expression of proliferating and contractile ASM cell receptors is limited so that the binding rate saturates with the combined density of proliferating or contractile ASM and active TGF-*β*. This leads to the following system of ODEs describing the rate of change in density of the two ASM sub-populations, ECM and concentration of active TGF-*β*:

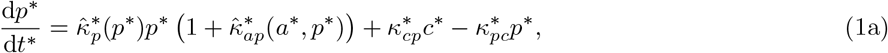

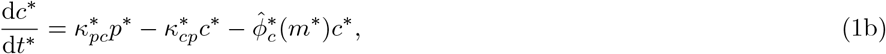

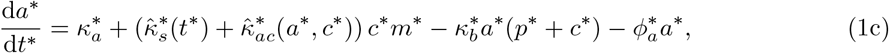

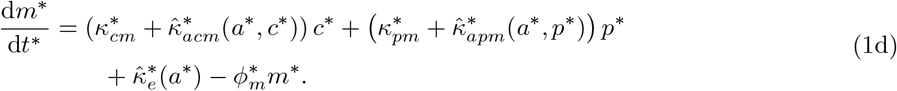

**Figure 1:**
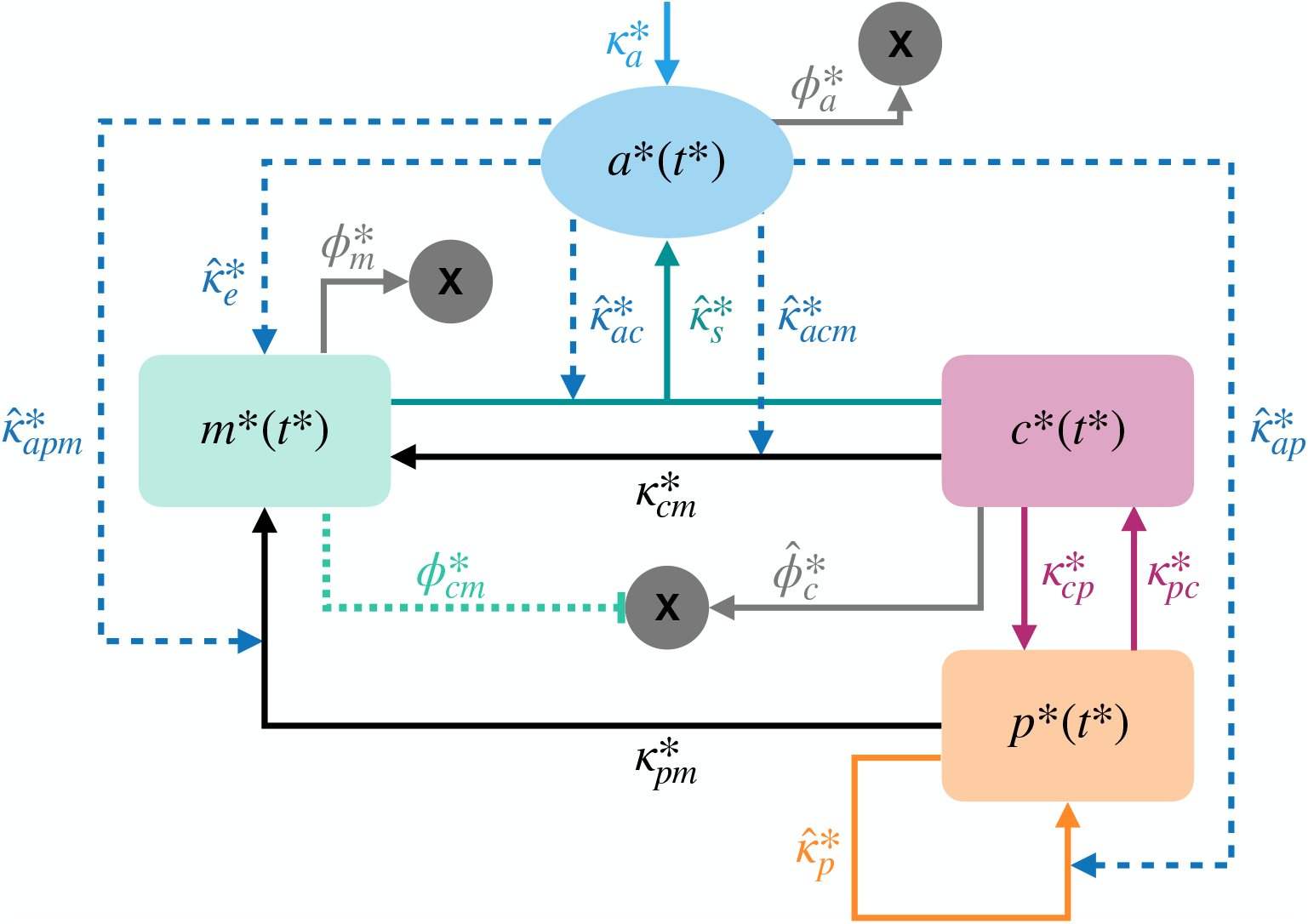
Schematic diagram detailing the interactions between the key model components: proliferating ASM, *p*\*(*t*\*), contractile ASM, *c*\*(*t*\*), ECM, *m*\*(*t*\*) and active TGF-*β, a*\*(*t*\*), at rates *κ* and *ϕ* as listed, explained and non-dimensionalised in Tables 1–3 in Appendix A. Dashed blue lines represent TGF-*β* signalling pathways and the grey circles, labelled X, represent degradation of the respective components.

Following Chernyavsky et al. (2014), we assume that phenotype switching may occur between these two ASM sub-populations. As described in Section 1, the roles of the proliferating ASM and contractile ASM differ; for simplicity, we assume the contractile ASM phenotype is exclusively responsible for contraction, whereas the proliferative phenotype is exclusively responsible for ASM growth. The rate constant 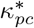 represents phenotype switching from the proliferative to the contractile state; 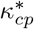 denotes switching in the reverse direction. We assume that the proliferation rate, 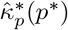, is of logistic form

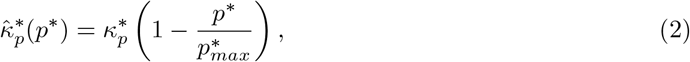

thereby implicitly accounting for a spatial constraint, with 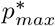 defining the carrying capacity of the proliferating ASM cell density. ASM proliferation is thought to increase with asthma severity (Johnson et al., 2001; Trian et al., 2007; Pelaia et al., 2008; Januskevicius et al., 2016) and correspondingly, the rate constant 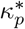, defining the rate of ASM proliferation, may be altered to reflect the severity of asthma.

The rate function 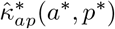 represents active TGF-*β*-induced proliferation and acts in addition to baseline proliferation 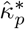, while 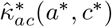 represents the rate of activation of additional TGF-*β* as a result of TGF-*β*-induced ASM contraction. The rate functions 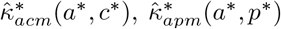 and 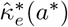 represent the rate of secretion of ECM by the contractile ASM, proliferating ASM and other surrounding cells, respectively, that is induced by active TGF-*β* signalling pathways. In addition, baseline secretion of ECM (independent of active TGF-*β* concentration) occurs at rates 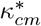 and 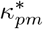 by the contractile and proliferative ASM cells, respectively. It is assumed that the density of proliferating and contractile ASM is not changed as a result of secretion of ECM.

Following the approach of Baker et al. (2017), the rates for TGF-*β*-induced ASM proliferation, 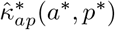, ECM secretion, 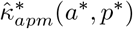, active TGF-*β*-induced ASM contraction, 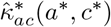, and ECM secretion, 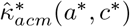 take the following form

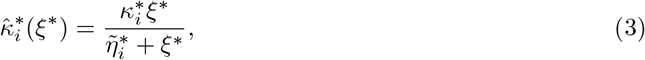

wherein the index *i* denotes each of the four rates *i* ∈ {*ap, apm, ac, acm*}, and the argument *ξ*\* represents *a*\**p*\* or *a*\**c*\*, respectively. The dependence on the product of the variables in each case reflects the initiation of these processes by binding of TGF-*β* to receptors expressed by proliferative or contractile ASM cells, respectively. We note that we consider first order saturation of these Hill functions, and those that follow, for simplicity.

We assume that the receptors expressed by other surrounding cells (excluding ASM cells) are in abundance so that the rate for active TGF-*β*-induced ECM secretion from external sources, 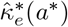, is

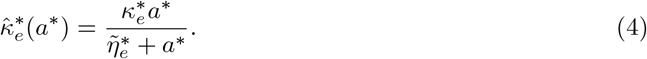

ECM presence has been shown to aid the survival of contractile ASM (Freyer et al., 2001). We therefore assume that the apoptosis rate of contractile ASM depends on the ECM density such that

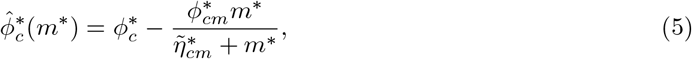

where, in the absence of ECM-aided survival, apoptosis of contractile ASM occurs at a maximum rate 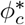. In order to ensure that the presence of ECM does not directly increase the density of ASM, yet reduces the ASM apoptosis rate to a minimum as the further addition of ECM no longer significantly aids ASM survival, we enforce 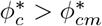. ECM degrades at rate 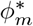.

Recall that ASM cell contraction mechanically activates latent TGF-*β* (Tatler and Jenkins, 2012), by unfolding the latent complex that is anchored to the ECM and that encapsulates latent TGF-*β*. Hence, both the contractile ASM and ECM are required for the mechanical activation of TGF-*β* (Tatler et al., 2011). This is reflected by the 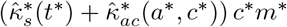 term in (1c). Latent TGF-*β* is found in abundance (Annes et al., 2003) and therefore is not a limiting factor in the mechanical activation step. Activation of latent TGF-*β* in response to an external stimulus, 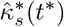, represents exposure to an irritant and mimics an asthmatic exacerbation. Experimentally, the stimulus may represent the administration of a contractile agonist such as methacholine (MCh) that is known to cause contraction of the ASM in lung-slice models (Tan and Sanderson, 2014). Following Chernyavsky et al. (2014), we model this as a series of Gaussian stimuli, given by

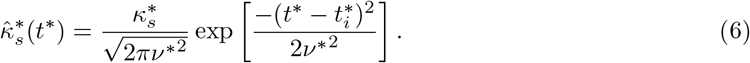

The stimulation occurs at times 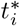 for a duration *ν* and with amplitude 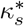. Subsequent active TGF-*β* signalling pathways drive further mechanical activation of latent TGF-*β* via ASM contraction, which is assumed to occur at rate 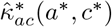, as in (3). Active TGF-*β* is a contractile agonist and therefore generates a positive feedback loop of further TGF-*β* activation. Basal activation of latent TGF-*β* via alternative sources is assumed to occur at a constant rate 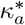.

In addition to the natural degradation of active TGF-*β*, 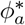, we must account for removal of active TGF-*β* through binding to the receptors expressed on both the proliferative and contractile ASM cells. Hence, we include the removal term

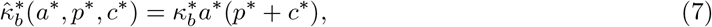

that describes the rate of change in concentration of active TGF-*β*.

The initial densities of ASM and ECM, and the concentration of active TGF-*β* are denoted by

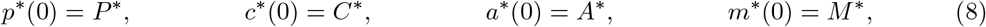

where *P*\*, *C*\*, *A*\*, *M*\* > 0.

### 2.1 Parameter choices

There is a lack of detailed experimental data to constrain the large number of parameters in our proposed model (1). For example, the study of Bai et al. (2000) quantifies long-term ASM mass accumulation in vivo to elucidate the contribution of age and disease duration on remodelling; however, observed increases may arise due to a variety of processes, posing problems in employing this data to quantify the relevant rate in (1). Conversely, quantitative data obtained from in vitro experiments do not necessarily reflect accurately corresponding processes in vivo. Therefore, we follow Baker et al. (2017) and choose an illustrative parameter set whereby comparable processes have similar rates, and demonstrate model behaviour by varying key parameters from these baseline values. Where possible, we determine estimates for the order of magnitude of rates from Tatler (2016) and the wider literature. The rate of natural degradation of active TGF-*β*, 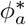, is one of the few known rates within the model and occurs on the order of minutes (Wakefield et al., 1990; Makinde et al., 2007). Baseline parameter values are given in Table 4 in Appendix A.

### 2.2 Non-dimensionalisation

We non-dimensionalise time, *t*\*, relative to the rate of natural degradation of active TGF-*β*, 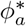, (as it is one of the few parameter values we are most confident of) such that

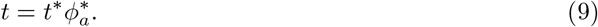

The density of proliferating ASM, *p*\*, contractile ASM, *c*\*, ECM, *m*\* and concentration of active TGF-*β, a*\*, are scaled with reference quantities, 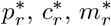 and 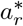, respectively, to obtain

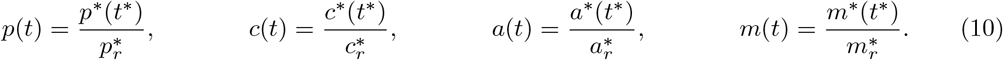

In view of (9), (10), the dimensionless version of the ODE model (1) is given by

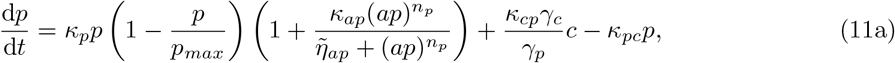

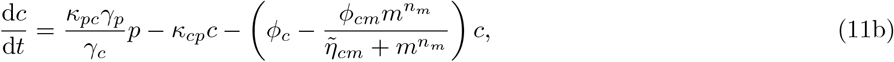

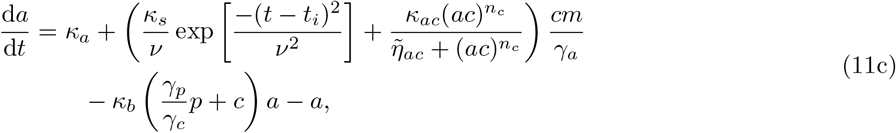

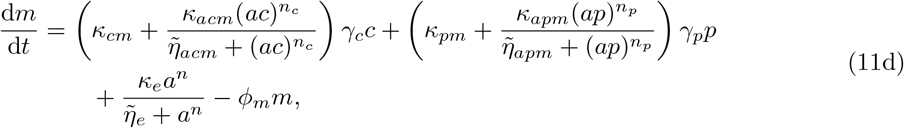

wherein the dimensionless parameters

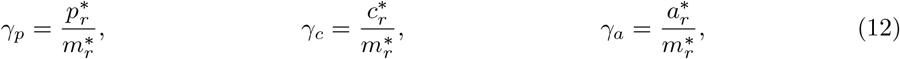

represent the ratio of the reference quantities of proliferating ASM, 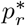, contractile ASM, 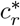, and active TGF-*β*, 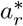, to that of ECM, 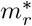, respectively. The non-dimensionalisation of the remaining parameters in the model (11) are provided in Tables 1–3 in Appendix A.

**Table 1:** Table of parameters in the ODE model (1), including the dimension, biological description and the corresponding dimensionless parameters in the dimensionless model (11).

| Parameter | Dimension | Description | Dimensionless |
| --- | --- | --- | --- |
| $p_r^*$ | $\frac{N}{L^3}$ | Reference number of proliferating ASM cells per unit volume of tissue. | N/A |
| $c_r^*$ | $\frac{N}{L^3}$ | Reference number of contractile ASM cells per unit volume of tissue. | N/A |
| $a_r^*$ | $\frac{N}{L^3}$ | Reference number of active TGF-β molecules per unit volume of tissue. | N/A |
| $m_r^*$ | $\frac{N}{L^3}$ | Reference number of ECM proteins per unit volume of tissue. | N/A |
| $P^*$ | $\frac{N}{L^3}$ | Initial number of proliferating ASM cells per unit volume of tissue. | $P = \frac{P^*}{p_r^*}$ |
| $C^*$ | $\frac{N}{L^3}$ | Initial number of contractile ASM cells per unit volume of tissue. | $C = \frac{C^*}{c_r^*}$ |
| $A^*$ | $\frac{N}{L^3}$ | Initial number of active TGF-β molecules per unit volume of tissue. | $A = \frac{A^*}{a_r^*}$ |
| $M^*$ | $\frac{N}{L^3}$ | Initial number of ECM proteins per unit volume of tissue. | $M = \frac{M^*}{m_r^*}$ |
| $p_{max}^*$ | $\frac{N}{L^3}$ | Maximum number of proliferating ASM cells per unit volume of tissue. | $p_{max} = \frac{p_{max}^*}{p_r^*}$ |
| $\gamma_a$ | 1 | Ratio of reference TGF-β concentration and reference ECM density. | $\gamma_a = \frac{a_r^*}{m_r^*}$ |
| $\gamma_p$ | 1 | Ratio of reference proliferating ASM and reference ECM density. | $\gamma_p = \frac{p_r^*}{m_r^*}$ |
| $\gamma_c$ | 1 | Ratio of reference contractile ASM and reference ECM density. | $\gamma_c = \frac{c_r^*}{m_r^*}$ |
| $t_i^*$ | T | Time of stimulus. | $t_i = \phi_a^* t_i^*$ |
| $\nu^*$ | T | Duration of stimulus. | $\nu = \sqrt{2} \phi_a^* \nu^*$ |

**Table 2:** Table of parameters in the ODE model (1), including the dimension, biological description and the corresponding dimensionless parameters in the dimensionless model 11.

| Parameter | Dimension | Description | Dimensionless |
| --- | --- | --- | --- |
| $\phi_a^*$ | $\frac{1}{T}$ | Rate of active TGF-β degradation. | $1 = \frac{\phi_a^*}{\phi_a^*}$ |
| $\phi_c^*$ | $\frac{1}{T}$ | Rate of contractile ASM apoptosis. | $\phi_c = \frac{\phi_c^*}{\phi_a^*}$ |
| $\phi_{cm}^*$ | $\frac{1}{T}$ | Rate of ECM aided contractile ASM survival. | $\phi_{cm} = \frac{\phi_{cm}^*}{\phi_a^*}$ |
| $\phi_m^*$ | $\frac{1}{T}$ | Rate of ECM degradation. | $\phi_m = \frac{\phi_m^*}{\phi_a^*}$ |
| $\kappa_p^*$ | $\frac{1}{T}$ | Rate of ASM proliferation. | $\kappa_p = \frac{\kappa_p^*}{\phi_a^*}$ |
| $\kappa_{ap}^*$ | 1 | Rate of TGF-β-induced ASM proliferation. | $\kappa_{ap} = \kappa_{ap}^*$ |
| $\kappa_{cp}^*$ | $\frac{1}{T}$ | Rate of switch from contractile ASM phenotype to proliferative ASM phenotype. | $\kappa_{cp} = \frac{\kappa_{cp}^*}{\phi_a^*}$ |
| $\kappa_{pc}^*$ | $\frac{1}{T}$ | Rate of switch from proliferative ASM phenotype to contractile ASM phenotype. | $\kappa_{pc} = \frac{\kappa_{pc}^*}{\phi_a^*}$ |
| $\kappa_a^*$ | $\frac{N}{L^3 T}$ | Rate of basal TGF-β activation. | $\kappa_a = \frac{\kappa_a^*}{\phi_a^* a_r^*}$ |
| $\kappa_s^*$ | $\frac{L^3}{N}$ | Rate of stimulus-induced TGF-β mechanical activation. | $\kappa_s = \frac{\kappa_s^* c_r^*}{\sqrt{\pi}}$ |
| $\kappa_{ac}^*$ | $\frac{L^3}{NT}$ | Rate of TGF-β-induced TGF-β mechanical activation. | $\kappa_{ac} = \frac{\kappa_{ac}^* c_r^*}{\phi_a^*}$ |
| $\kappa_{cm}^*$ | $\frac{1}{T}$ | Rate of ECM secretion from contractile ASM. | $\kappa_{cm} = \frac{\kappa_{cm}^*}{\phi_a^*}$ |
| $\kappa_{acm}^*$ | $\frac{1}{T}$ | Rate of TGF-β-induced ECM secretion from contractile ASM. | $\kappa_{acm} = \frac{\kappa_{acm}^*}{\phi_a^*}$ |
| $\kappa_{pm}^*$ | $\frac{1}{T}$ | Rate of ECM secretion from proliferating ASM. | $\kappa_{pm} = \frac{\kappa_{pm}^*}{\phi_a^*}$ |
| $\kappa_{apm}^*$ | $\frac{1}{T}$ | Rate of TGF-β-induced ECM secretion from proliferating ASM. | $\kappa_{apm} = \frac{\kappa_{apm}^*}{\phi_a^*}$ |

**Table 3:** Table of parameters in the ODE model (1), including the dimension, biological description and the corresponding dimensionless parameters in the dimensionless model 11.

| Parameter | Dimension | Description | Dimensionless |
| --- | --- | --- | --- |
| $\kappa_e^*$ | $\frac{N}{L^3 T}$ | Rate of TGF-β-induced ECM deposition from alternative sources. | $\kappa_e = \frac{\kappa_e^*}{\phi_a^* m_r^*}$ |
| $\kappa_b^*$ | $\frac{L^3}{NT}$ | Rate of Active TGF-β binding to a receptor presented on proliferating or contractile ASM. | $\kappa_b = \frac{\kappa_b^* c_r^*}{\phi_a^*}$ |
| $\tilde{\eta}_{ap}^*$ | $(\frac{N^2}{L^6})^{n_p}$ | Disassociation constant. | $\tilde{\eta}_{ap} = \frac{\tilde{\eta}_{ap}^*}{(a_r^* p_r^*)^{n_p}}$ |
| $\tilde{\eta}_{apm}^*$ | $(\frac{N^2}{L^6})^{n_p}$ | Disassociation constant. | $\tilde{\eta}_{apm} = \frac{\tilde{\eta}_{apm}^*}{(a_r^* p_r^*)^{n_p}}$ |
| $\tilde{\eta}_{ac}^*$ | $(\frac{N^2}{L^6})^{n_c}$ | Disassociation constant. | $\tilde{\eta}_{ac} = \frac{\tilde{\eta}_{ac}^*}{(a_r^* c_r^*)^{n_c}}$ |
| $\tilde{\eta}_{acm}^*$ | $(\frac{N^2}{L^6})^{n_c}$ | Disassociation constant. | $\tilde{\eta}_{acm} = \frac{\tilde{\eta}_{acm}^*}{(a_r^* c_r^*)^{n_c}}$ |
| $\tilde{\eta}_{cm}^*$ | $(\frac{N}{L^3})^{n_m}$ | Disassociation constant. | $\tilde{\eta}_{cm} = \frac{\tilde{\eta}_{cm}^*}{(m_r^*)^{n_m}}$ |
| $\tilde{\eta}_e^*$ | $(\frac{N}{L^3})^n$ | Disassociation constant. | $\tilde{\eta}_e = \frac{\tilde{\eta}_e^*}{(a_r^*)^n}$ |

Dimensionless initial data is given by:

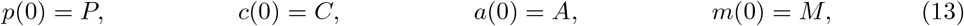

where,

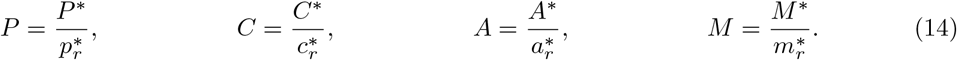

## 3 Asymptotic reduction

From Tatler (2016), we identify two timescales in the model: a fast timescale associated with rapid signalling pathways, and a slow scale describing gradually evolving downstream effects. The basal non-mechanical activation of TGF-*β, κ*_*a*_, ECM-aided survival of contractile ASM, *ϕ*_*cm*_, ECM secretion from the contractile ASM, *κ*_*cm*_ and *κ*_*acm*_, and ECM secretion from other surrounding cells, *κ*_*e*_, act on a slow timescale of hours to days, compared to the remaining rates within the systems that act over the order of minutes. Therefore, we introduce a small parameter, 0 *< ε* ≪ 1, and scale *κ*_*a*_, *ϕ*_*cm*_, *κ*_*cm*_, *κ*_*acm*_ and *κ*_*e*_ such that

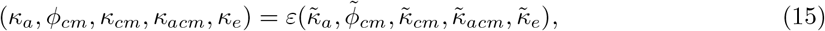

and expand each of the dependent variables *ψ* ∈ {*p, c, a, m*} in an asymptotic series such that

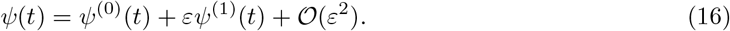

Substituting the expansions (16) into the model (11), we obtain at leading order:

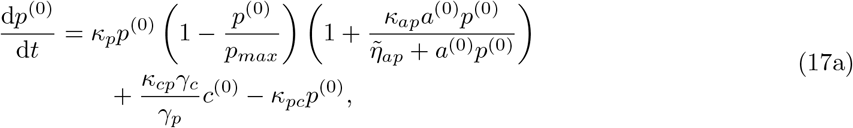

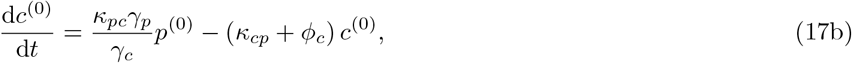

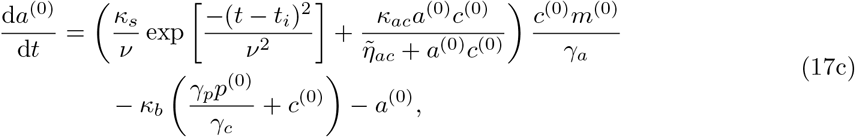

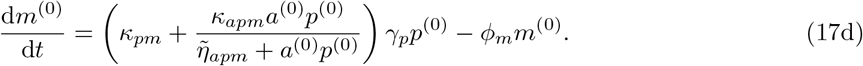

The above leading-order reduction will be used subsequently to identify steady states semi-analytically that will guide our stability analyses of the fully nonlinear model (11).

## 4 Stability analysis

In the following, we identify the steady states of the reduced system (17) and determine the stability of the trivial steady state analytically (Section 4.1). Thereby, we identify key parameters controlling the transition from trivial to biologically-relevant states. In Section 4.2 we perform a numerical bifucation analysis of the full nonlinear model (11) numerically, concentrating on those parameters identified in Section 4.1.

### 4.1 Asymptotically reduced model

Here, we exploit the reduction (17) to obtain semi-analytically the steady states of the system and to interrogate their stability. In the absence of stimulation, i.e. *κ*_*s*_ = 0, we find that there are four possible non-negative steady states. Specifically, equations (17) admit the trivial steady state

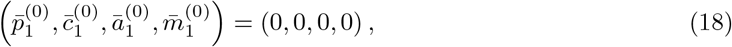

and a second state in which only the concentration of active TGF-*β* is zero,

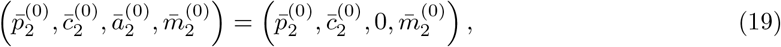

where,

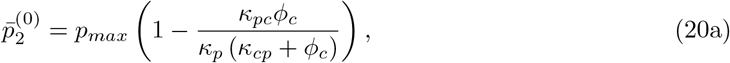

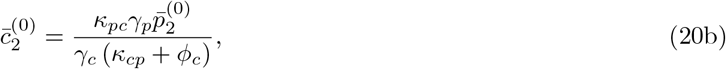

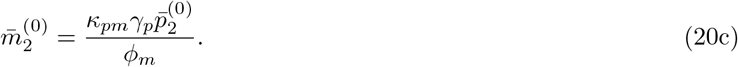

The overbars in (18)–(20) and hereafter indicate the steady state of each variable and subscripts denote each of the four steady state sets. The steady state (19) is positive and thus, biologically relevant, for

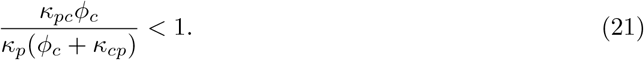

A further two positive steady states are given by the intersection of the nullclines of (17):

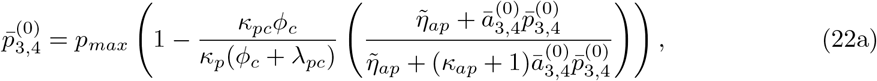

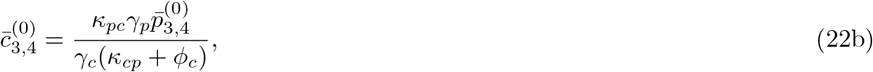

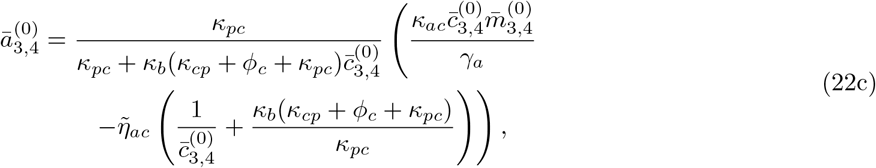

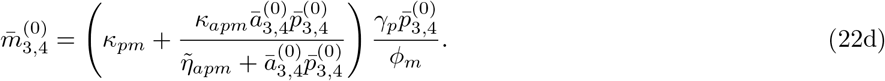

Explicit solutions to (22) are not possible and hence we obtain them numerically for a range of suitable parameter estimates.

The stability of the leading order steady states is determined by the eigenvalues of the corresponding Jacobian matrix of (17). Below, we consider the trivial steady state (18), for which eigenvalues can be obtained analytically. The Jacobian matrix for the reduced system (17) evaluated at (18), **J**_0_, is given by

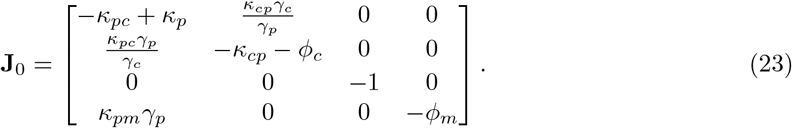

Correspondingly, the eigenvalues, Λ_*j*_ for *j* ∈ {1, …, 4}, of **J**_0_ are given by

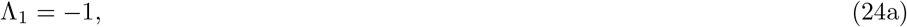

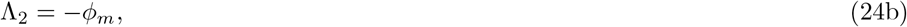

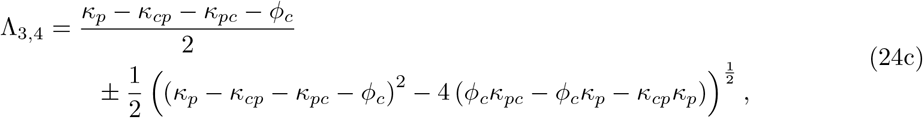

The eigenvalue Λ_3_ is positive for

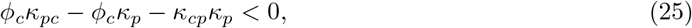

and thus, under this restriction, the trivial state (18) is unstable. The inequality (25) rearranges to (21) and hence, this instability condition coincides with the existence of the biologically relevant steady state (19). Accordingly, we have located a transcritical bifurcation. We note that the rate of ASM phenotype switching from the proliferative to contractile state, *κ*_*pc*_, and the rate of ASM proliferation, *κ*_*p*_, are of similar magnitude and thus (21) and (25) are satisfied for biologically relevant parameters.

### 4.2 Full model stability analysis

Guided by our findings in the reduced model (17) in Section 4.1, here we carry out a bifurcation analysis of the full model (11). The dimensionless parameters representing the rate of ASM proliferation, *κ*_*p*_, phenotype switching from the proliferative to contractile phenotype, *κ*_*pc*_ and contractile ASM degradation, *ϕ*_*c*_, (all relative to the rate of active TGF-*β* degradation) feature in the reduced restriction (21) that determines the key transition from the trivial steady state to a positive, biologically relevant, steady state. We therefore investigate the stability and location of the steady states of the full model (11) as functions of these parameters (Fig. 2). All bifurcation diagrams are produced using the dynamical systems software XPPAUT.

**Figure 2:**
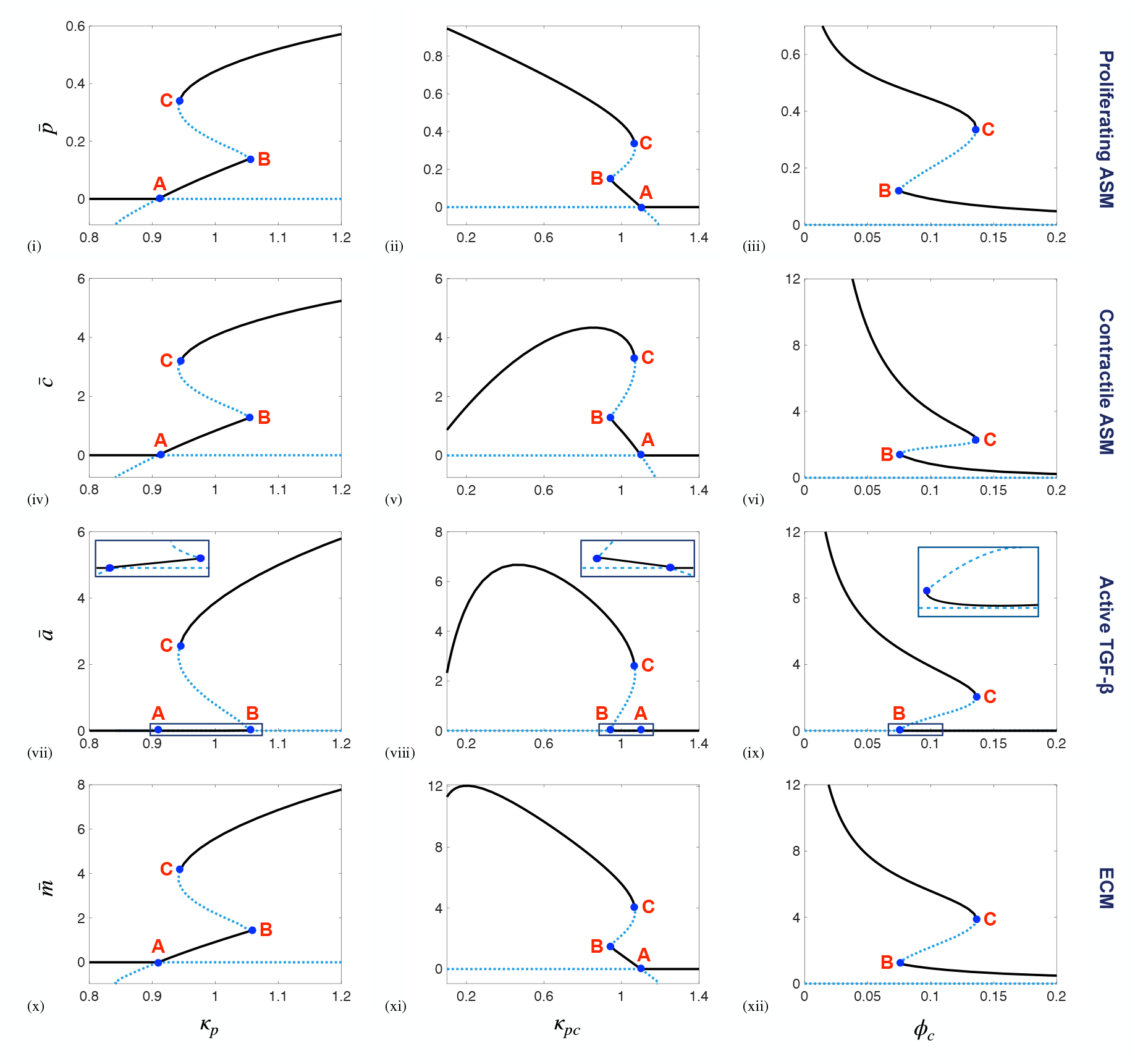
Bifurcation diagrams for *κ*_*p*_, *κ*_*pc*_, *ϕ*_*c*_ (first, second and third column, respectively) in the model (11). The parameter values are baseline values provided in Table 4 in Appendix A.

As previously identified in the reduced model (17), we observe regimes where there are up to four possible non-negative steady states. Within the chosen parameter ranges of *κ*_*p*_ and *κ*_*pc*_, we observe three distinct bifurcation points: a transcritical bifurcation, a change in stability of the positive branch and finally, another change in stability that gives rise to a bistable regime (points A, B, C, respectively in the first two columns of Fig. 2). In the reduced model (17), these observations correspond to the transcritical bifurcation in the zero steady state (18) as the non-trivial steady state (19) becomes positive and stable by satisfying the parameter restriction (21). The other two leading order steady states correspond to the upper and lower bound of the bistable region. Furthermore, we have verified that for parameter values where the asymptotic approximation is valid, the leading order values for the bifurcation points are within 5% of the values determined from the full model.

As outlined in Section 1, there is a correlation in the severity of asthma and the amount of active TGF-*β* present in the airways (Chen and Khalil, 2006; Al Alawi et al., 2014). Within the bistable parameter regime, we hence associate the stable steady state where the concentration of active TGF-*β* remains low (on the lower stable branched labelled AB in Fig. 2) with a healthy homeostatic state that maintains normal ASM and ECM accumulation, and that on the upper branch (created at C), where the concentration of active TGF-*β* is chronically elevated, with a diseased state that results in excessive ASM and ECM accumulation. The unstable branch between the two acts as a transition threshold leading to irreversible airway alterations and asthma development.

For values of *κ*_*p*_ below the location of the transcritical bifurcation point (A), there are no biologically relevant steady states in the system (11) (see Fig. 2, first column), suggesting that the proliferation rate must be large enough to allow for maintainence of the ASM and ECM. For increasingly large *κ*_*p*_, the bistable regime is surpassed (point B, Fig. 2, first column) and the elevated positive steady state becomes the only biologically relevant steady state. Intuitively, this steady state rises with increasing rate of ASM proliferation due to the accumulation of ASM which subsequently increases ECM secretion and causes further activation of TGF-*β*.

On the other hand, there are no biologically relevant steady states in the system (11) for values of *κ*_*pc*_ above the transcritical bifurcation point (Fig. 2, second column). This indicates that the proliferative-to-contractile switching rate must be low enough that there are sufficiently many proliferative ASM cells to maintain ASM mass. The stable population of proliferating ASM increases with decreasing switching rate; in contrast, contractile ASM, ECM and active TGF-*β* initially increase with *κ*_*pc*_, before subsequently decreasing (Fig. 2 (v,viii,xi)). It is interesting to note in passing that the maxima in these variables occur at different values of *κ*_*pc*_. In sum, these results indicate that there is an optimal switching rate where there is sufficient proliferative ASM to maintain ASM and ECM density, yet allowing for a sufficiently large contractile ASM population needed to mechanically activate TGF-*β* (Fig. 2 (ii), (v)).

The two bifurcation points at which there is a change in direction and stability of the positive stable branch also appear as we vary the rate of degradation of the contractile ASM, *ϕ*_*c*_ (Fig. 2, third column; labelled B, C). For large values of *ϕ*_*c*_ there is a transcritical bifurcation, as observed for *κ*_*p*_ and *κ*_*pc*_; however, the location of this is not visible and we have restricted the range of *ϕ*_*c*_ that is displayed in the third column of Fig. 2 in order to study the bistable regime of interest closely. Here, the lower region of the stable branch reduces towards the unstable steady state as *ϕ*_*c*_ increases. In contrast, the positive steady-state values all increase dramatically for small values of *ϕ*_*c*_, highlighting the key role of contractile ASM apoptosis in preventing excessive ASM growth, ECM secretion and TGF-*β* activation.

In Figure 3 we track the motion of the bifurcation points A, B, C shown in Fig. 2 in two dimensional parameter space to show the creation and destruction of the bistable regime. Within the studied parameter range, we see that the transcritical bifurcation point (A; Fig. 2) does not exist for all values of *ϕ*_*c*_, *κ*_*p*_, *κ*_*pc*_ (Fig. 3(i, iii)), but is present for all pairs (*κ*_*p*_, *κ*_*pc*_) (Fig. 3(ii)). In particular, for values of *κ*_*p*_ *>* 1 and for values of *κ*_*pc*_ *<* 1 only the bistable regime exists (Fig. 3 (i, iii)). Thus, a balance is required between the rate of proliferation and apoptosis of the ASM in order to permit a bistable regime that exhibits a state of relatively low concentration of active TGF-*β*. As proliferation increases, so must the degradation of ASM to regulate the increased ASM mass, subsequent ECM secretion and TGF-*β* activation.

**Figure 3:**
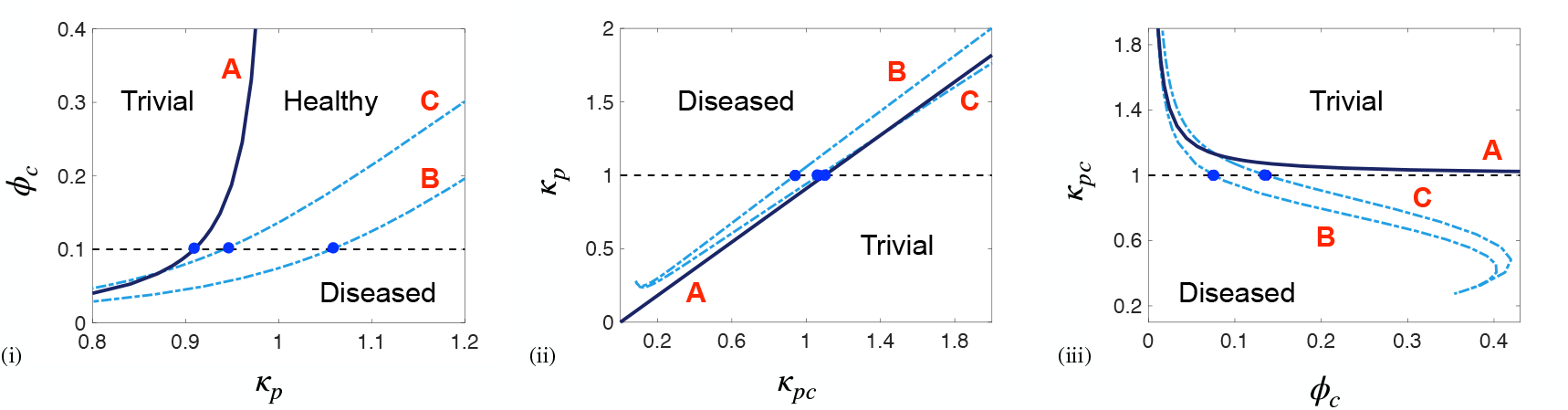
Two parameter bifurcation diagrams for *κ*_*p*_, *κ*_*pc*_, *ϕ*_*c*_ in the model (11). Solid lines track the transcritical branching point, blue dashed lines track the limit points defining the bistable regime, black dashed lines indicate a slice corresponding to Fig. 2 and the blue dots represent the corresponding bifurcation points. The parameter values are baseline values provided in Table 4 in Appendix A.

For very small values of *κ*_*pc*_, we see that there are no bifurcation points for any value of *ϕ*_*c*_, reflecting the fact that in this case, there is so little contractile ASM, due to insignificant amounts being switched from the proliferative population, that the degradation of what remains does not make a significant difference to the overall dynamics of the system. Similarly, for very small values of *κ*_*p*_, we see that not all values of *κ*_*pc*_ permit a bistable regime and the transcritical bifurcation is the only bifurcation point in the system (Fig. 3(ii)).Biologically, the amount of proliferating ASM that may be exchanged to the cont.ractile state is is considerably reduced, due to the decline in the production rate. Hence, any minor contribution from the proliferative population is insignificant, irrespective of the switch rate, and does not alter the overall system dynamics.

## 5 Time-course simulation results

The bistable region we have identified above allows for a switch between a healthy homeostasis, in which the concentration of active TGF-*β* remains very small, and a diseased state of a chronically elevated concentration of active TGF-*β*. The intermediate unstable branch acts as a transition threshold. In this section we demonstrate how this manifests in simulated disease development and treatment.

### 5.1 Effect of asthmatic exacerbations

In this section, we demonstrate that a series of stimuli, 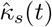 in (6), can drive the system out of the healthy homeostatic state and into the diseased state. Following the approach of Chernyavsky et al. (2014), we simulate multiple asthmatic exacerbations at time points, *t*_*i*_, via the dimensionless full model (11). We initialise the model (11) with the healthy homeostatic state and show the possible agonist-driven increase in the activation of TGF-*β* and the downstream consequences that may lead to airway remodelling in asthma. All simulated parameter values, selected within the bistable regime, are provided in the relevant figure captions and in Table 5 in Appendix A.

**Table 4:** Table of dimensionless baseline parameter values for the ODE model (11).

| Parameter | Baseline | Parameter | Baseline | Parameter | Baseline |
| --- | --- | --- | --- | --- | --- |
| $\phi_c$ | 0.1 | $\gamma_p$ | 1 | $\tilde{\eta}_{ap}$ | 1 |
| $\phi_m$ | 0.01 | $\gamma_a$ | 0.01 | $\tilde{\eta}_{apm}$ | 10 |
| $\phi_{cm}$ | 0.001 | $\gamma_c$ | 1 | $\tilde{\eta}_{ac}$ | 1 |
| $\kappa_p$ | 1 | $\kappa_{ap}$ | 1 | $\tilde{\eta}_{acm}$ | 1 |
| $\kappa_{cp}$ | 0.01 | $\kappa_{pc}$ | 1 | $\tilde{\eta}_{cm}$ | 1 |
| $\kappa_a$ | 0.001 | $\kappa_{ac}$ | 0.01 | $\tilde{\eta}_e$ | 1 |
| $\kappa_{apm}$ | 0.1 | $\kappa_{pm}$ | 0.1 | $p_{max}$ | 1 |
| $\kappa_{acm}$ | 0.001 | $\kappa_{cm}$ | 0.0001 | | |
| $\kappa_e$ | 0.001 | $\kappa_b$ | 1 | | |

**Table 5:** Table of dimensionless parameter values for the ODE model (11) used to produce Fig. 4 – 7. The remaining parameter values are baseline values, as provided in Table 4.

| **Fig. 4** | **Parameter** |  |  |
| --- | --- | --- | --- |
| | $t_i$ | $\nu$ | $\kappa_s$ |
| (i) | 50:40:250 | 5 | 0.1 |
| (ii) | 50:40:450 | 5 | 0.1 |
| (iii) | 50:40:650 | 5 | 0.1 |
|  |  |  |  |
| **Fig. 5** |  |  |  |
| (i) | 50:70:750 | 5 | 0.1 |
| (ii) | 50:50:550 | 5 | 0.1 |
| (iii) | 50:30:350 | 5 | 0.1 |
|  |  |  |  |
| **Fig. 6** |  |  |  |
| (i) | 50:50:550 | 20 | 0.1 |
| (ii) | 50:50:550 | 10 | 0.1 |
| (iii) | 50:50:550 | 1 | 0.1 |
|  |  |  |  |
| **Fig. 7** |  |  |  |
| (i) | 50:50:300 | 5 | 0.1 |
| (ii) | 50:50:300 | 5 | 0.15 |
| (iii) | 50:50:300 | 5 | 0.17 |

**Table 6:**
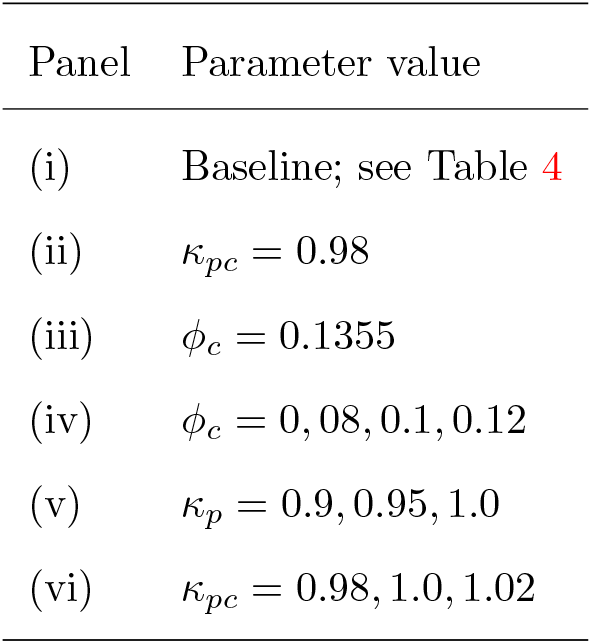
Table of dimensionless parameter values for the ODE model (11) used to produce Fig. 8. The remaining parameter values are baseline values, as provided in Table 4.

The effect of increasing the frequency of recurrent challenges of contractile agonist with fixed amplitude is illustrated in Figure 4. For a low number of challenges, we observe a transient increase in active TGF-*β* concentration followed by a return to the healthy homeostatic state (Fig. 4 (i)). As the number of exposures increases, the active TGF-*β* concentration continues to climb (see, e.g., Fig. 4 (ii)) leading to ASM and ECM mass increases and further activation of TGF-*β* in a positive feedback loop. In this instance, the threshold for which the system can return to the healthy state is surpassed and the irreversible changes to the overall dynamics lead to the diseased state.

**Figure 4:**
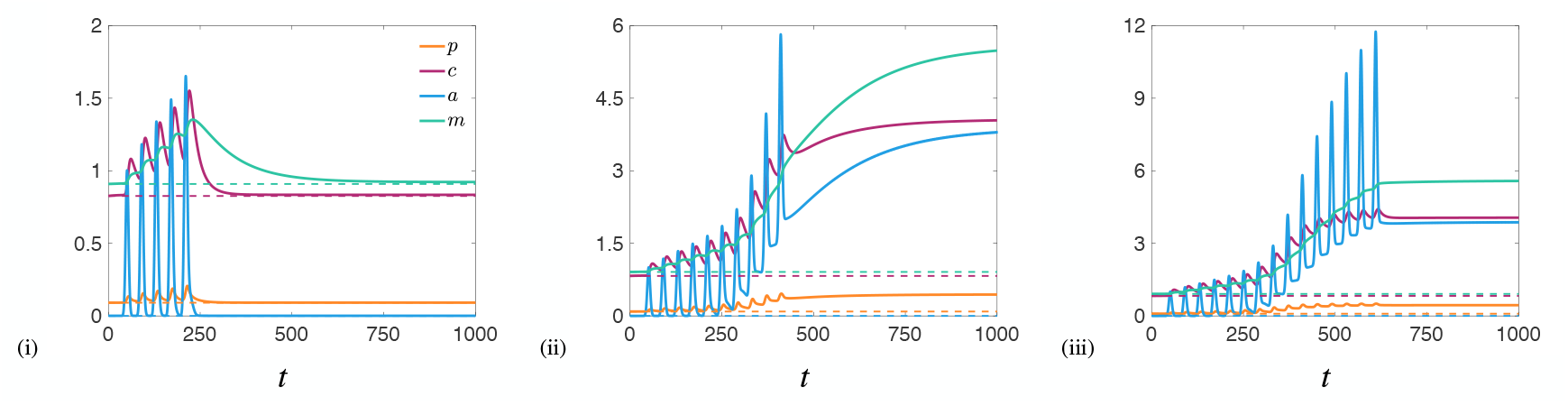
Results from the model (11) to illustrate the effect of increasing the number of stimuli (6) driving TGF-*β* activation on the density of airway wall components. In panels (i)–(iii), the number of exposures are 5, 10 and 15. Dashed lines correspond to the healthy homeostatic state, at which we initialise the system. Parameter values, selected within the bistable regime, are provided in Table 5 in Appendix A.

The effect of reducing the period between exposures for a fixed number of recurrent stimuli is shown in Fig. 5. When the stimulation occurs at a low frequency there is sufficient recovery time between exposures for the system to return the healthy homeostatic state (Fig. 5 (i), (ii)). As the frequency increases, adequate clearance of active TGF-*β* is hindered (Fig. 5 (iii)) and thus the mechanotransductive feedback loop of further TGF-*β* activation drives the system towards the diseased state.

**Figure 5:**
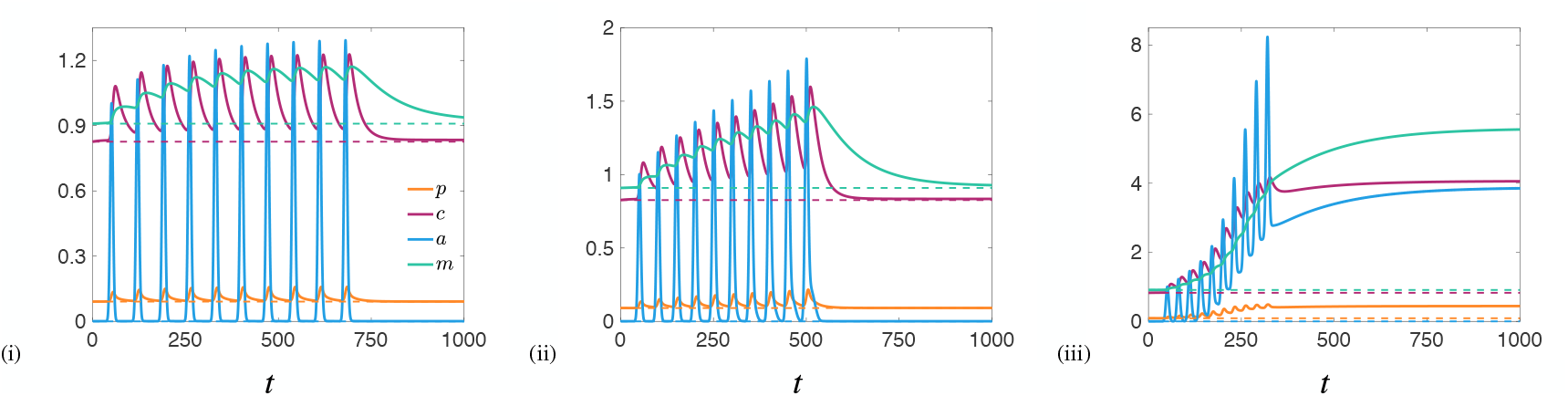
Results from the model (11) to illustrate the effect of increasing the frequency of stimuli (6) driving TGF-*β* activation on the density of airway wall components. The frequency of exposures increase between plots from (i)–(iii). Dashed lines correspond to the healthy homeostatic state initial condition. Parameter values, selected within the bistable regime, are provided in Table 5 in Appendix A.

As is the case for increasing stimulus frequency (Fig. 5), we find that reducing the rate of active TGF-*β* clearance (following exposure to a fixed number of recurrent stimuli, occurring at the same frequency) leads to the re-establishment of a chronically elevated TGF-*β* concentration (Fig. 6). The persistent presence of active TGF-*β* due to reduced clearance affects the downstream dynamics and ASM and ECM continues to accumulate following the final exacerbation (Fig. 6 (i, ii)). Conversely, sufficiently fast clearance enables complete removal of active TGF-*β* after each challenge so that, while ASM and ECM periodically rise and fall, the increase is insufficient to drive the system into the diseased state and so on cessation of the challenges it returns to the healthy homeostatic state (Fig. 6 (iii)). We note that despite clearance, the small accumulations of ASM and ECM within the challenge period leads to an increasing activation of TGF-*β* with subsequent challenges, as was the case in the previous cases of the diseased state (see Fig. 6 (iii), 4 and 5); here the TGF-*β* level eventually reaches a consistent maximum following exposure to the stimulant.

**Figure 6:**
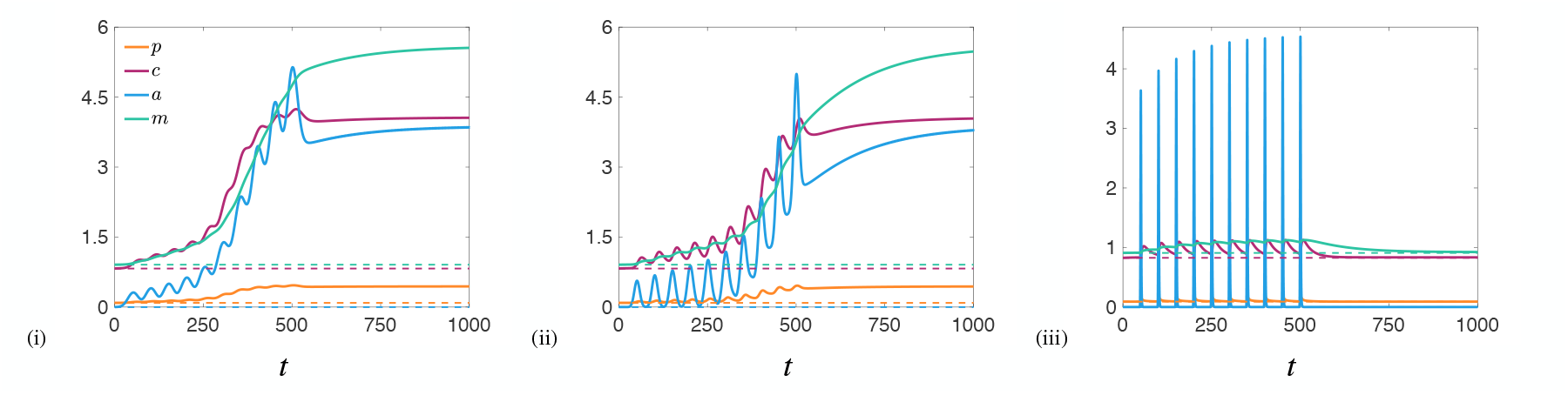
Results from the model (11) to illustrate the effect of increasing the clearance of active TGF-*β, ν* in (6), following stimuli, on the density of airway wall components. The value of *ν* increases between plots from (i)–(iii). Dashed lines correspond to the healthy homeostatic state initial condition. Parameter values, selected within the bistable regime, are provided in Table 5 in Appendix A.

The effect of increasing the exposure amplitude, *κ*_*s*_, for a given number of recurrent stimuli, occurring at fixed frequency and clearance rate, is illustrated in Fig. 7. As expected, we see that for relatively low amplitude stimulation, the long-term dynamics are unaffected by the temporary rise in ASM, ECM and active TGF-*β* accumulation (Fig. 7 (i)). Increasing *κ*_*s*_ (Fig. 7 (ii)), we observe a slight delay in the return from the temporary peak to the healthy steady state. This is highlighted by the accumulation of ASM and active TGF-*β* (Fig. 7 (ii)). Further increase gives rise to the disease state (Fig. 7 (iii)). We find that the model is very sensitive to the strength of the stimulant and that very little change is required in order to significantly alter the downstream dynamics. Here, the percentage increase in *κ*_*s*_ (the parameter representing the strength of the exacerbation) that is required to exhibit a significant change in the long-term behaviour is at least an order of magnitude smaller than in previous parameter explorations.

**Figure 7:**
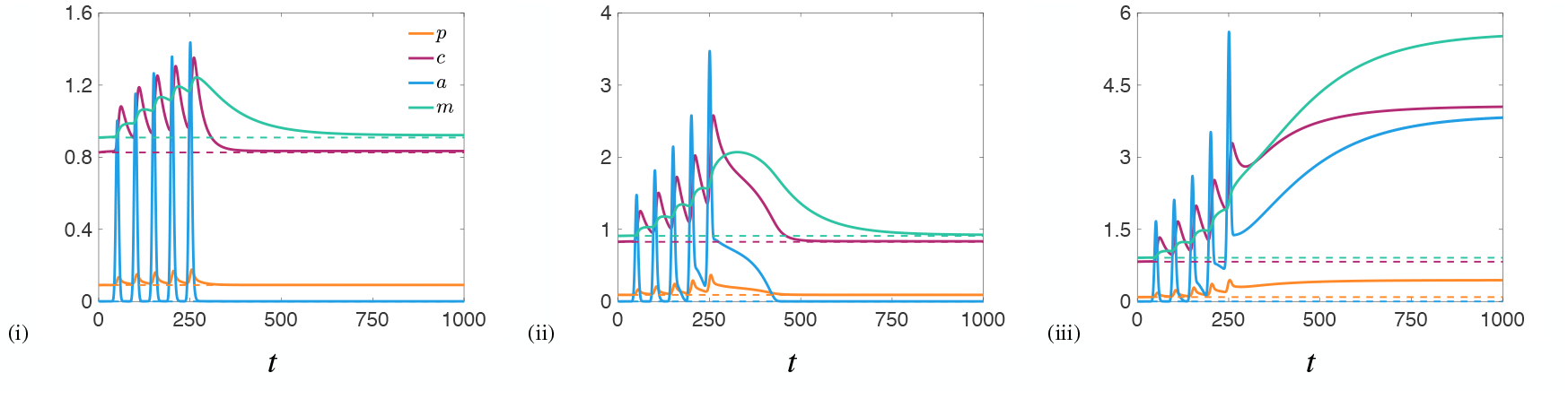
Results from the model (11) to illustrate the effect of increasing the strength of stimuli, *κ*_*s*_ in (6), that drives TGF-*β* activation, on the density of airway wall components. The value of *κ*_*s*_ increases between plots from (i)–(iii). Dashed lines correspond to the healthy homeostatic state initial condition. Remaining parameter values, selected within the bistable regime, are provided in Table 5 in Appendix A.

### 5.2 Effect of ASM reduction via bronchial thermoplasty

Here we initialise the system at the diseased steady state, depicted (e.g.) by the points on the curve labelled B in Figure (3), and simulate the temporal response to a sudden decrease in contractile and proliferative ASM density as a percentage of their initial value (mimicking the effect of bronchial thermoplasty) for a range of percentage removal and parameter values *κ*_*p*_, *κ*_*pc*_ and *ϕ*_*c*_. Example time-courses are shown in Figures 8(i–iii). Thereby, we determine the minimum ablation required for the system to leave the diseased state and return to the (non-trivial) healthy steady state, consequently reversing the effects of asthmatic airway remodelling for a given model parameter regime. These results are summarised in Figure 8(iv–vi), highlighting that the ablation threshold shows strong and non-trivial dependence on the parameter regime. For example, in the case of an ASM population that is highly proliferative (large *κ*_*p*_), has suppressed apoptosis (low *ϕ*_*c*_), or slow switching from proliferative to contractile phenotype (small *κ*_*pc*_), the model predicts that thermoplasty is not appropriate treatment, requiring 100% ASM removal for efficacy (Fig. 8(iv–vi), blue lines). However, due to the nonlinear relationship between ablation threshold and parameter values, a relatively small reduction in ASM proliferation results in a significant decrease in this threhold (Fig. 8(iv), blue line); in combination with increases in phenotypic switching or apoptosis, further dramatic reductions are achievable (Fig. 8(v,vi), orange lines). Importantly, we observe regions in parameter space in which the model predicts successful treatment under physiologically realistic levels of ablation. These model predictions thereby point to possible biomarkers under which thermoplasty is likely to be successful.

**Figure 8:**
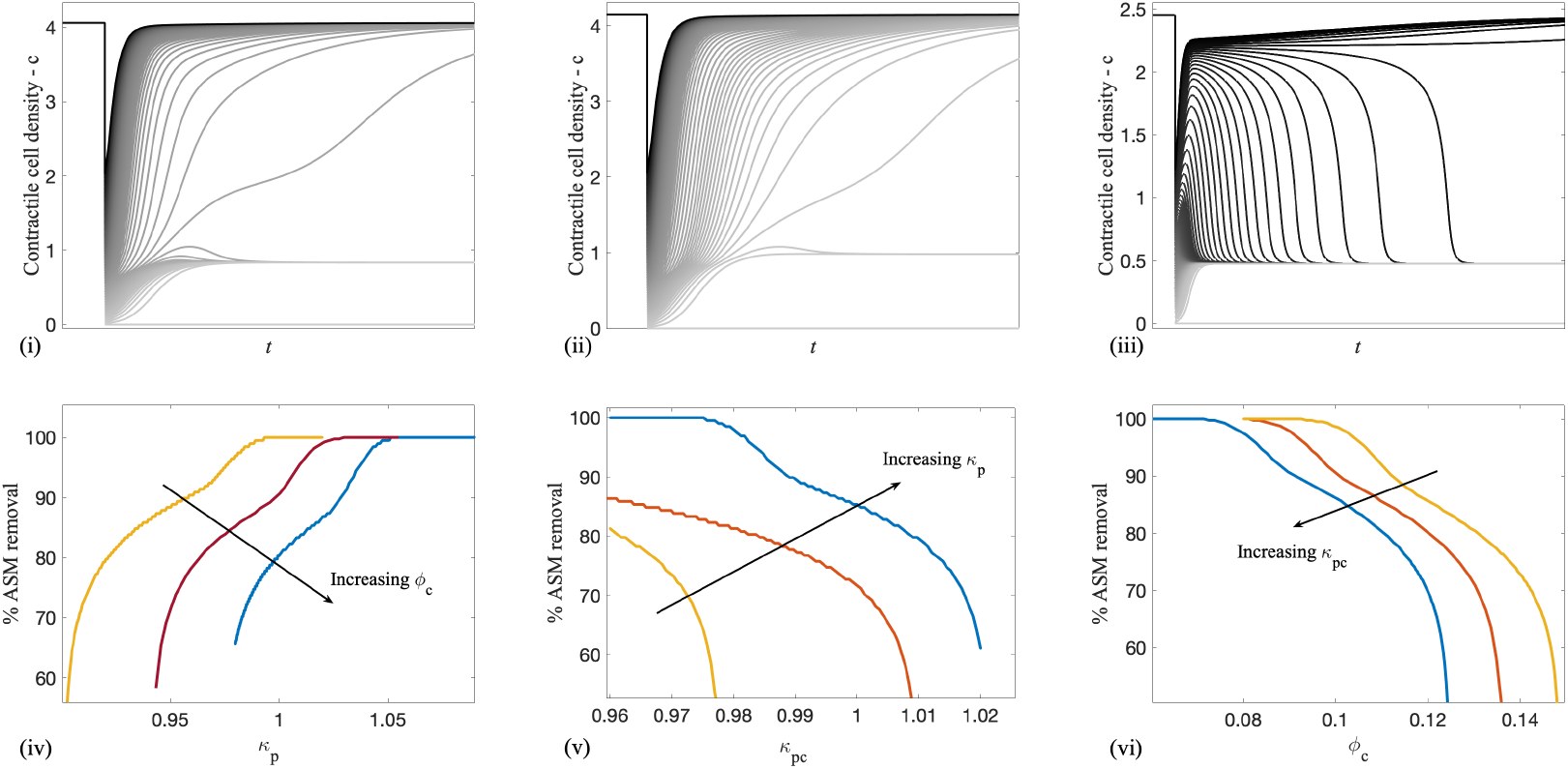
Results from the model (11) to illustrate the response to bronchial thermoplasty under different parameter choices. (i)–(iii): time course simulations showing the response of the contractile cell density to simulated ablation at (i) baseline parameter values, (ii) slightly reduced *κ*_*pc*_, (iii) slightly increased *ϕ*_*c*_. ASM removal (both *c* and *p*) varies from 50% to 100% in steps of 0.5%. (iv)–(vi): The minimum ablation required to drive the system from the diseased state to the non-trivial healthy steady state, as a function of the parameters *κ*_*p*_, *κ*_*pc*_ and *ϕ*_*c*_. Parameter values, selected within the bistable regime, are provided in Table 6 in Appendix A.

## 6 Discussion

In this study, we have presented a new model to describe the interaction of ASM, ECM and TGF-*β* activation and downstream conseqeunces, on the change in density of the airway wall constituents. Estimates of the timescales of these interactions allowed us to obtain a reduced model that permitted semi-analytical progress.Thereby, we determined the important parameters that dominate the transition from homeostasis to disease, on which we then focus in a numerical bifurcation analysis of the nonlinear model. These results expose a bistable region in parameter space allowing for the co-existence of both a steady state maintaining a relatively low concentration of active TGF-*β*, representing healthy homeostasis, and a higher concentration of active TGF-*β* with accompanying increased ASM and ECM density, representing a diseased state. We find that the rate of ASM proliferation, degradation and proliferative-to-contractile ASM phenotype switching dominate the long-term dynamics of the system. The ASM is the link between each of these factors and thus, our findings suggest that the ASM, in both a contractile and proliferative state, is the key component within the airway wall responsible for maintaining growth, ECM secretion and TGF-*β* mechanical activation. We have shown that a balance is required between these three important roles to allow for the overall system to exhibit a bistable regime. Our model does not currently account for changes in ASM-ECM adhesion, but the bistable system dynamics associated with these short time-scale mechanical behaviours (see e.g. the earlier-cited biomechanical studies (Irons et al., 2018)) may play a further role in activation of TGF-*β* and should be considered in future studies.

Simulations of external stimuli mimicking a series of asthmatic exacerbations highlight the importance of the number, the strength, and frequency of exacerbation occurrence and the clearance of active TGF-*β*. We observed that both an increased frequency of exposures to a stimulant and a reduced TGF-*β* clearance rate increases the accumulation in ASM and ECM and alters the steady state of the system in the long-term. This may be apparent in patients who experience regular exposures to an allergen and as a result, develop long-term irreversible airway remodelling. We have shown that allowing adequate recovery time (e.g. avoiding allergens) or improving recovery time (i.e. reduced frequency and sufficient active TGF-*β* clearance) between successive exacerbations returns the system to a healthy homeostatic state. Therefore, in theory, increasing the clearance of active TGF-*β* could be a possible method of asthma management or treatment.

We investigate the temporal dynamics of airway constituents and TGF-*β* following a reduction in ASM content mimicking the implementation of bronchial thermoplasty in severe asthmatics. Numerical simulations showed that a certain amount of damage to the ASM is needed to ensure that the constituent densities are driven down to the healthy steady state and that this threshold depends on the underlying parameters. Identifying such a threshold is important because it is clear from previous studies that the technique is not able to remove all the ASM, and that in fact the amount of damage as measured via biopsies in clinical trials may be closer to 50% (Chernyavsky et al., 2018). Our finding additionally suggests that patient response to treatment could depend not only the acute damage to the ASM immediately after thermoplasty but also on longer term response which will depend on their specific disease phenotype (Papakonstantinou et al., 2021).

Our findings in Section 5.1 are consistent with Chernyavsky et al. (2014) and Hill et al. (2018), who likewise explore airway remodelling in response to transient inflammatory or contractile agonist challenges. Similarly, we find that when the clearance of active TGF-*β* is inefficient (i.e. the inflammation resolution speed is slow), a small number of exacerbation events may lead to significant airway remodelling that remains after the stimulant resides. Chernyavsky et al. (2014) consider remodelling due to the growth of the ASM alone (ASM hyperplasia) and found that the phenotype switching rate from the contractile to proliferative state is modulated by the inflammatory status (which changes in response to the stimulant). In contrast, we model the accumulation of ECM and active TGF-*β* accompanying ASM growth and consider a constant switching rate between the ASM sub-populations. In future work, we could consider a switch rate that is dependent upon the concentration of active TGF-*β*. Other considerations could include development towards a stochastic model and simulating likely outcomes due to exposures occurring at irregular intervals.

The morphoelastic model of Hill et al. (2018) reveals persistent contractile tone in asthmatics via either a mechanotransductive feedback loop, insufficient clearance of contractile agonists, or a combination of the two. Complementary to these findings, our ODE model uncovers a bistable regime and highlights a threshold concentration of active TGF-*β* that once surpassed, leads to irreversible airway remodelling. Motivated by these findings, in forthcoming work we will couple subcellular signalling dynamics to our previous nonlinear hyperelastic model of airways in precision-cut lung-slices (Pybus et al., 2021) to study mechanochemical mechanisms underpinning asthma characteristics in vivo.

## Acknowledgements

Dr Hannah J. Pybus was supported by the London Mathematical Society Early Career Fellowship, Grant number: ECF-1920-49, and acknowledges the generous support of the QJMAM Fund for Applied Mathematics for providing funding for a collaborative research visit related to this work thereafter.

We thank Dr Amanda L. Tatler (Division of Respiratory Medicine, University of Nottingham, UK) for assisting with biological matters and providing advice regarding suitable model parameters.

## A Model parameters

Tables of parameter values corresponding to the model (11) developed in Section 2.

## Notes

### Competing Interest Statement

The authors have declared no competing interest.

